# Natural recombination among Type I restriction-modification systems creates diverse genomic methylation patterns among *Xylella fastidiosa* strains

**DOI:** 10.1101/2022.09.01.506293

**Authors:** Michael L. O’Leary, Lindsey P. Burbank

## Abstract

*Xylella fastidiosa* is an important bacterial pathogen of plants causing high consequence diseases in agricultural crops around the world. Although as a species *X. fastidiosa* can infect an extremely broad range of host plants, significant variability exists between strains and subspecies groups in virulence on specific host plant species, and other traits such as growth habits. Natural competence and horizontal gene transfer are believed to occur frequently in *X. fastidiosa*, and likely influences the evolution of this pathogen. However, some *X. fastidiosa* strains are extremely difficult or impossible to manipulate genetically using standard transformation techniques. Several restriction-modification systems are encoded in the *X. fastidiosa* genome, including multiple Type I R-M systems that may influence horizontal gene transfer and recombination. In this study, several conserved Type I R-M systems were compared across 129 *X. fastidiosa* genome assemblies representing all known subspecies and 32 sequence types. Considerable allelic variation among strains was identified among the single specificity subunit (*hsdS*) of each Type I R-M system, with a unique *hsdS* allele profile generally conserved within a monophyletic cluster of strains. Inactivating mutations were identified in Type I R-M systems of specific strains, showing heterogeneity in the complement of functional Type I R-M systems across *X. fastidiosa*. Genomic DNA methylation patterns were characterized in 20 *X. fastidiosa* strains and associated with Type I R-M system allele profiles. Overall, this study describes epigenetic modifications in *X. fastidiosa* associated with functional Type I R-M systems and characterizes the diversity in these systems across *X. fastidiosa* lineages.

**Importance:** Economic impacts on agricultural production due to *X. fastidiosa* have been severe in the Americas, Europe, and parts of Asia. Despite a long history of research on this pathogen, certain fundamental questions regarding the biology, pathogenicity, and evolution of *X. fastidiosa* have still not been answered. Wide scale whole genome sequencing has begun to provide a more insight into *X. fastidiosa* genetic diversity and horizontal gene transfer but the mechanics of genomic recombination in natural settings and extent to which this directly influences bacterial phenotypes such as plant host range are not well understood. Genome methylation is an important factor in horizontal gene transfer and bacterial recombination that has not been comprehensively studied in *X. fastidiosa*. This study characterizes methylation associated with Type I restriction-modification systems across a wide range of *X. fastidiosa* strains and lays the groundwork for a better understanding of *X. fastidiosa* biology and evolution through epigenetics.

## Introduction

*Xylella fastidiosa* is an insect-transmitted xylem-limited bacterial plant pathogen capable of causing disease on numerous agricultural and ornamental host species (1). Globally, this pathogen has significant impacts on agricultural production, and is the subject of substantial quarantine regulation in international plant trade (EFSA 2019). Strains of *X. fastidiosa* can be separated into as many as five genetically distinct subspecies (i.e., *fastidiosa, morus, multiplex, pauca*, and *sandyi*), that are further categorized into sequence types (STs) according to a multi-locus sequence typing scheme (3). The host range of *X. fastidiosa* as a species is very broad, but the number of host plant species upon which a given ST of *X. fastidiosa* will cause severe disease appears to be limited (4), and recombination between strains may result in pathogenicity on novel hosts (5). To date there is very little understanding of the genetic determinants of plant host range in *X. fastidiosa*, and there is no way to effectively and specifically predict host range of newly identified strains based on DNA sequence information alone. Although *X. fastidiosa* is naturally competent and homologous recombination has been implicated in genetic diversity and evolution of this species (6–8), several aspects of the underlying mechanics of recombination are still unclear. In addition to natural transformation (9), *X. fastidiosa* can undergo conjugative DNA transfer (10), and prophage integration is also evident throughout the genome (11, 12). However, some strains of *X. fastidiosa* are much more difficult to transform then others, and there is variability in degree of natural competence between strains (13–15). Although restriction-modification systems have been implicated in *X. fastidiosa* transformation efficiency, only a few of these have been specifically characterized so far (16, 17).

Restriction-modification (R-M) systems are prokaryotic self-recognition systems with methyltransferase and restriction endonuclease activity that allow bacterial cells to differentiate between self and non-self DNA by modifying and/or degrading DNA at specific sequence motifs (18). R-M systems play a role in defense against phages or other invasive DNA elements, but also can influence strain evolution by limiting genetic exchange between strains with differing R-M system complements (18, 19). R-M system abundance typically correlates with genome size, with many species carrying approximately one R-M system per Mb. However, naturally competent bacteria and species with reduced genomes (i.e., < 2 Mb) can carry significantly more R-M systems per Mb. Four types of R-M systems have been described, and the methyltransferase activity of these systems commonly modifies either adenine residues to generate N^6^-methyl-adenine (6mA), or cytosine residues to generate N^5^-methyl-cytosine (5mC) or N^4^-methyl-cytosine (4mC) (20). Furthermore, epigenetic modifications made by R-M systems can also influence gene expression. Alteration in methylation targets, including through stochastic on/off switching of R-M systems (phase variation), can generate bacterial subpopulations with different phenotypes including altered virulence (20, 21).

Type I R-M systems consist of a protein complex with both methylation and restriction endonuclease activity targeting double-stranded DNA at a bipartite motif separated by a non-specific spacer. This R-M complex is formed by subunit components responsible for modification (HsdM), specificity (HsdS), and restriction activity (HsdR) (22). The specificity subunit, HsdS, is necessary for both modification and restriction activity, and contains two variable target recognition domains (TRDs) separated by a central conserved region (CCR). While *hsdM* and *hsdR* are highly conserved, *hsdS* alleles with differing TRDs can be associated with a single Type I R-M system across or within strains. The sequence specificity of an *hsdS* allele is largely dependent on the specific TRDs found on that allele. Each TRD in an *hsdS* allele (i.e., TRD-1 and TRD-2) recognizes a short nucleotide sequence, typically 3-5 bp, corresponding to one-half of the bipartite motif targeted by that allele. TRDs can be highly divergent, with those that recognize different nucleotide sequences typically sharing little to no nucleotide or amino acid sequence similarity. Recombination events between *hsdS* genes can generate new combinations of TRDs (i.e., new alleles) with novel nucleotide specificity, allowing Type I R-M systems to rapidly evolve to target new genomic sites. Finally, Type I R-M systems can be phase-variable by reversible mutation, typically either through the presence of multiple *hsdS* subunits within a system which exchange position or recombine to generate variation, or variation in length of simple sequence repeat (SSR) tracts within *hsdS* or *hsdM* components (20, 21, 23–25).

Previous analysis of *X. fastidiosa* strains identified several DNA methyltransferases and restriction endonucleases including Type I R-M systems, and the presence of subspecies and strain-specific R-M systems (11, 16, 26). Transformation efficiency of some *X. fastidiosa* strains is increased by inhibition or deletion of Type I R-M systems (17). Type I R-M systems in *X. fastidiosa* strains appear to be genetically variable, containing both RFLP-identified variation within the *hsdR* subunit component, recombination within *hsdM* and *hsdR* subunit components, inactivating mutations (i.e., frameshifts), or loss of *hsdM* and/or *hsdS* components in some Type I R-M systems (13, 26–28). Furthermore, a recent survey of R-M systems available in REBASE indicated specific *X. fastidiosa* strains contain Type I R-M system components with SSR tracts that could cause phase-variable expression (23). However, the full variation among Type I R-M systems in *X. fastidiosa* and their biological relevance is not known.

Here, we used a genomics approach to investigate Type I R-M systems present in *X. fastidiosa* strains across 112 previously released genome assemblies, 15 new genome assemblies, and two new hybrid assemblies of previously sequenced strains (Bakersfield-1 and Stag’s Leap). We identify three Type I R-M systems conserved in all 129 *X. fastidiosa* genome assemblies, including one with homology to a Type I R-M system present in genome assemblies of 31 *Xylella taiwanensis* strains, and one additional Type I R-M system found only in *X. fastidiosa* subsp. *multiplex* and *pauca* strains. We characterized the single *hsdS* gene associated with each Type I R-M system, identifying 44 unique TRDs arranged in combination to form at least 50 unique *hsdS* alleles across the four Type I R-M systems. We show that these alleles are arranged in 31 allele profiles that are each typically, but not always, limited to a single monophyletic strain cluster of *X. fastidiosa*, and identify mutations in specific strains that may disrupt these systems, including SSR length variation. Finally, we identified differential epigenetic motifs associated with activity of Type I R-M systems and other methyltransferases in 20 *X. fastidiosa* strains, including representatives from each *X. fastidiosa* subspecies.

## Results and Discussion

### Genome sequencing and assembly results

*X. fastidiosa* strains sequenced in this study have typical genome size and GC content, and frequently contain plasmids, consistent with previous results for subsp. *morus* and *multiplex* (Table 1) (29, 30). Riv13 contains a previously unidentified plasmid (pXF62-Riv13) with 98% identity to a plasmid found in subsp. *pauca* strain Hib4, pXF64-HB (CP009886.1). OC8, a subsp. *sandyi* strain isolated from olive in Orange County, California, USA, contains a 30 kb plasmid (pXF30-OC8) identical to that of subsp. *sandyi* strain Ann-1 (CP006697.1). A similar plasmid is also present in Oak 35874 (pXF26-Oak35874). Some variation in potentially erroneously truncated coding sequences (e.g., sequencing errors) was observed, although these values were relatively consistent within subsp. *fastidiosa* and subsp. *multiplex, morus*, and *sandyi* and may be typical of *X. fastidiosa* genome assemblies.

**Table 1.**
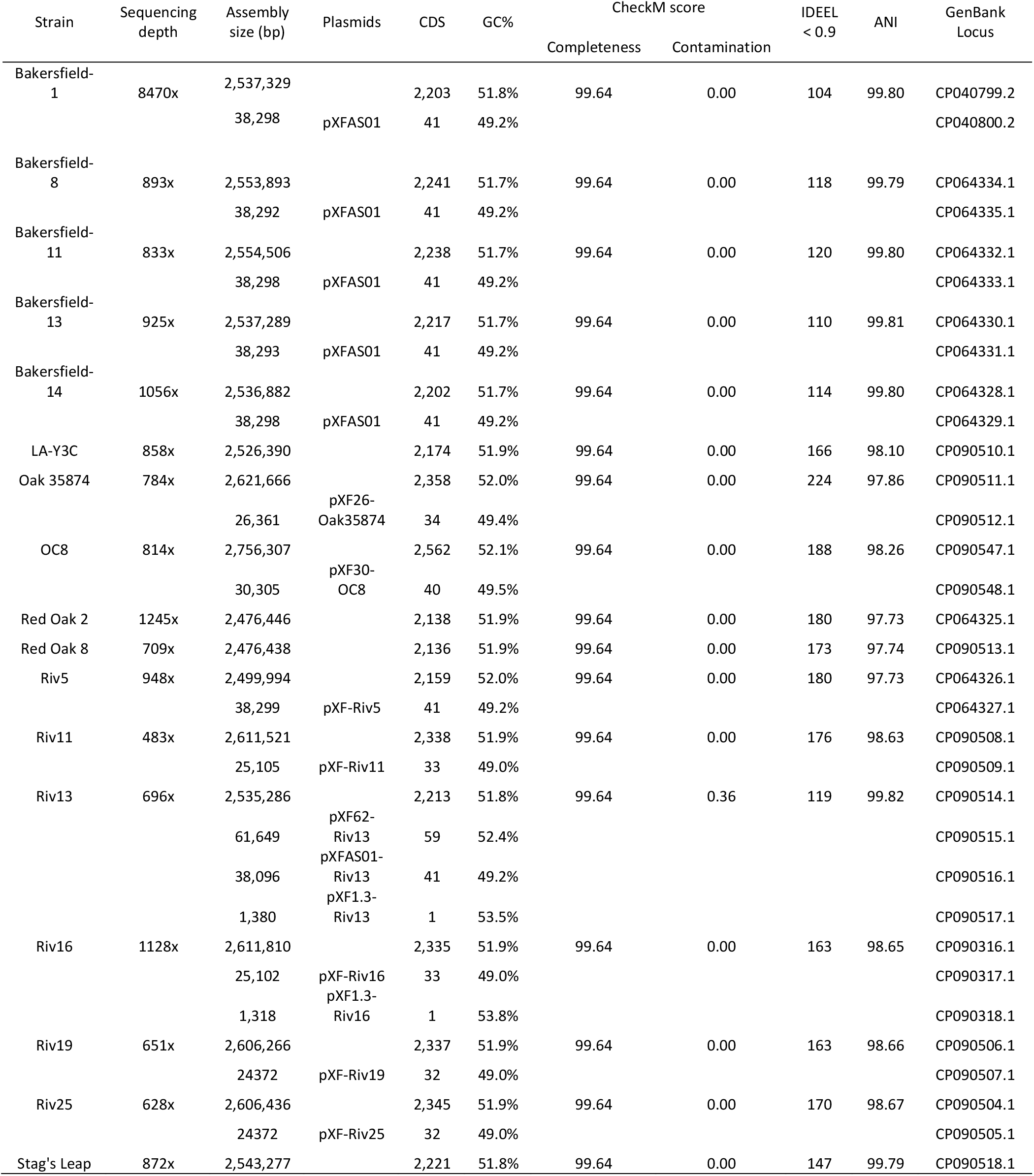
Strains sequenced in this study. ANI = Average nucleotide identity to *Xylella fastidiosa* type strain ATCC 35871. IDEEL < 0.9 = number of potentially truncated (e.g., frameshifted) CDS identified by IDEEL, relative to best UniProt TREMBL match.

### *Xylella fastidiosa* strains encode up to three Type I R-M systems within a conserved defense island

Comparison of *hsdM* and *hsdR* gene sequences from 129 *X. fastidiosa* strains identified four distinct Type I R-M systems, designated XfaI-XfaIV (Figure 1A and 1B). Three of these systems (XfaI, XfaII, and XfaIV) are present in all 129 *X. fastidiosa* genome assemblies. Strains of subsp. *multiplex* and *pauca* also encode a fourth Type I R-M system, XfaIII. Only one *hsdS* subunit was identified per system, located between the *hsdM* and *hsdR* components. Although *hsdM* and *hsdR* gene sequences of each system are highly conserved across strains and subspecies, *hsdS* gene sequences are substantially more variable (Figure 1A). This variation indicates the presence of distinct *hsdS* alleles, likely with different DNA recognition motifs, among these conserved systems. In a small number of strains (e.g., *X. fastidiosa* subsp. *pauca* strain 11399, subsp. *morus* strains), XfaI and/or XfaII appear to be incomplete (Table S1) (13).

**Figure 1.**
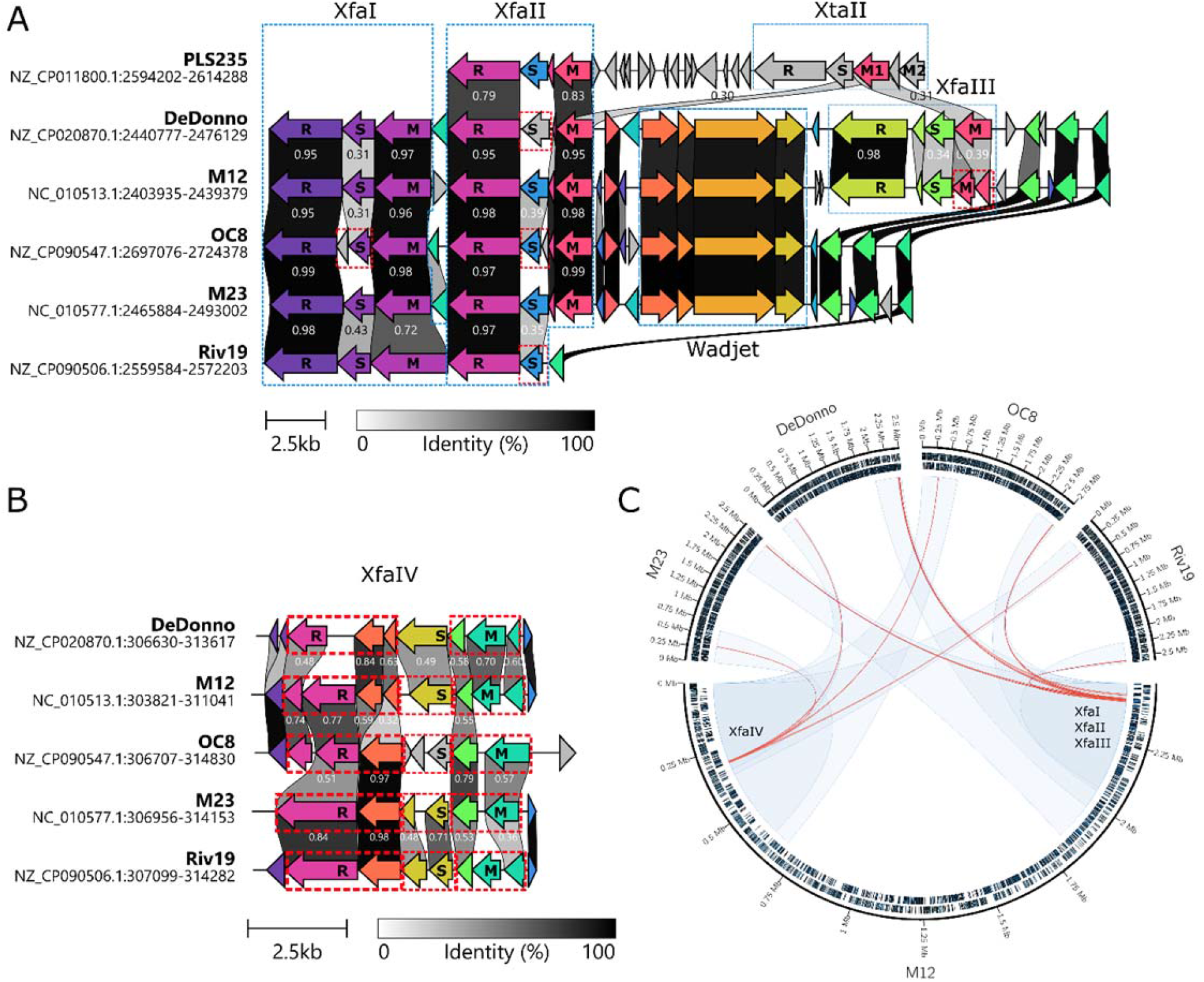
Alignment and genomic location of Type I Restriction-Modification systems identified in representative *Xylella fastidiosa* strains. A. Alignment of a putative defense island containing XfaI, XfaII, and XfaIII in strains where these systems are present. Distinct Type I R-M systems and the Wadjet system are indicated in blue boxes, with R-M system subunits labeled R = restriction subunit *hsdR*, M = methyltransferase subunit *hsdM*, S = specificity subunit *hsdS*. Red boxes indicate a fragmented gene; percent identity of translated coding sequences are given. B. Alignment of regions containing XfaIV, showing this system has acquired multiple inactivating mutations in each strain. C. Genomic locations of Type I R-M systems in representative *Xylella* strains in comparison to reference strain M12. Locally colinear blocks, identified with progressiveMauve, which contain type I restriction-modification systems are indicated by blue ribbons. Red lines indicate homology among Type I restriction-modification systems.

In addition to high nucleotide identity, these systems are positionally homologous in complete genome assemblies. Pairwise genome alignments of representative strains from each subspecies identified two locally collinear blocks in each strain which contain Type I R-M systems (Figure 1C). The first locally colinear block contains XfaIV, while the second contains XfaI, XfaII, and XfaIII. XfaI and XfaII are adjacent in all strains, separated by conserved sequence, with XfaIII located ∼10kb upstream on the negative strand only in subsp. *multiplex* and *pauca* strains. Analysis of this intervening sequence indicates it encodes a Wadjet type I defense system (31), which is present in all strains except subsp. *morus* strains. Wadjet type I defense systems are expected to provide defense against single-strand DNA, such as some phages or plasmid conjugation. In *X. fastidiosa*, this system may serve a complementary function to the Type I R-M systems, which recognize double-stranded DNA. Wadjet systems are typically seen sporadically within a species, so the presence of one in nearly every currently available genome sequence of *X. fastidiosa* is unusual.

Although XfaI and XfaII are present in all examined *X. fastidiosa* strains, some strains have acquired inactivating mutations in or deletions of individual subunits of one or more Type I R-M systems. Inactivation can be limited to specific strains within a subspecies, such as strain M12 (subsp. *multiplex*), where the *hsdM* subunit of XfaIII is fragmented. In all examined subsp. *morus* strains however, a deletion eliminates the *hsdM* subunit of XfaII, but not *hsdS or hsdR*. Most notable is XfaIV, where every examined strain from all subspecies has accrued at least one mutation that introduces frameshifts or early stop codons in one or more components of this system (Figure 1B). It is not clear why this system is retained in all *X. fastidiosa* strains, although similar retention of a nonfunctional Type I R-M system was observed in *Staphylococcus epidermidis* (32).

The presence of 3 conserved Type I R-M systems in all *X. fastidiosa* subspecies and a fourth system in subsp. *multiplex* and *pauca*, along with the conserved genomic location of these systems, suggests that they were acquired by a common ancestor of *X. fastidiosa* before these lineages diverged, rather than horizontally transferred between strains. Taken together, this evidence suggests 1) XfaI, XfaII, and XfaIV were acquired by a common ancestor of these *X. fastidiosa* strains, 2) specific lineages either gained or lost XfaIII after subspecies diverged, and 3) XfaI, XfaII, XfaIII, and the Wadjet system may constitute a defense island common to known *X. fastidiosa* strains.

### *Xylella taiwanensis* shares one of its three Type I R-M systems with *X. fastidiosa*

The closely related bacterial species *Xylella taiwanensis*, which causes similar leaf scorch diseases, is the only other species characterized in the *Xylella* genus and to date has only been found Taiwan (33). The genome of *X. taiwanensis* also encodes three Type I R-M systems, one of which is homologous to XfaII (Figure 1A, Table 2). The two other systems in *X. taiwanensis* are not found in *X. fastidiosa*, and are designated XtaI and XtaII respectively. In *X. taiwanensis* strain PLS235, XfaII is located ∼7 kb downstream of XtaII (Figure 1A). Unlike the Type I R-M systems identified in *X. fastidiosa*, XtaII encodes two distinct methyltransferases. The presence of a second methyltransferase suggests this system may be able to make both m6A and m4C modifications (24). Only one *hsdS* gene was identified per Type I-RM system in *X. taiwanensis*. Homology between XfaII in *X. fastidiosa* and *X. taiwanensis* suggests a common origin for this system. It’s possible that XfaII was acquired by a common ancestor of these two species, or perhaps transferred between them. Unlike *X. fastidiosa*, the 31 strains of *X. taiwanensis* examined here do not have *hsdS* allele variation within either XfaII, XtaI, or XtaII (Table 2, Table S2). However, it is plausible that the genetic diversity among *X. taiwanensis strains* sampled thus far is insufficient to identify *hsdS* allele variation.

**Table 2.**
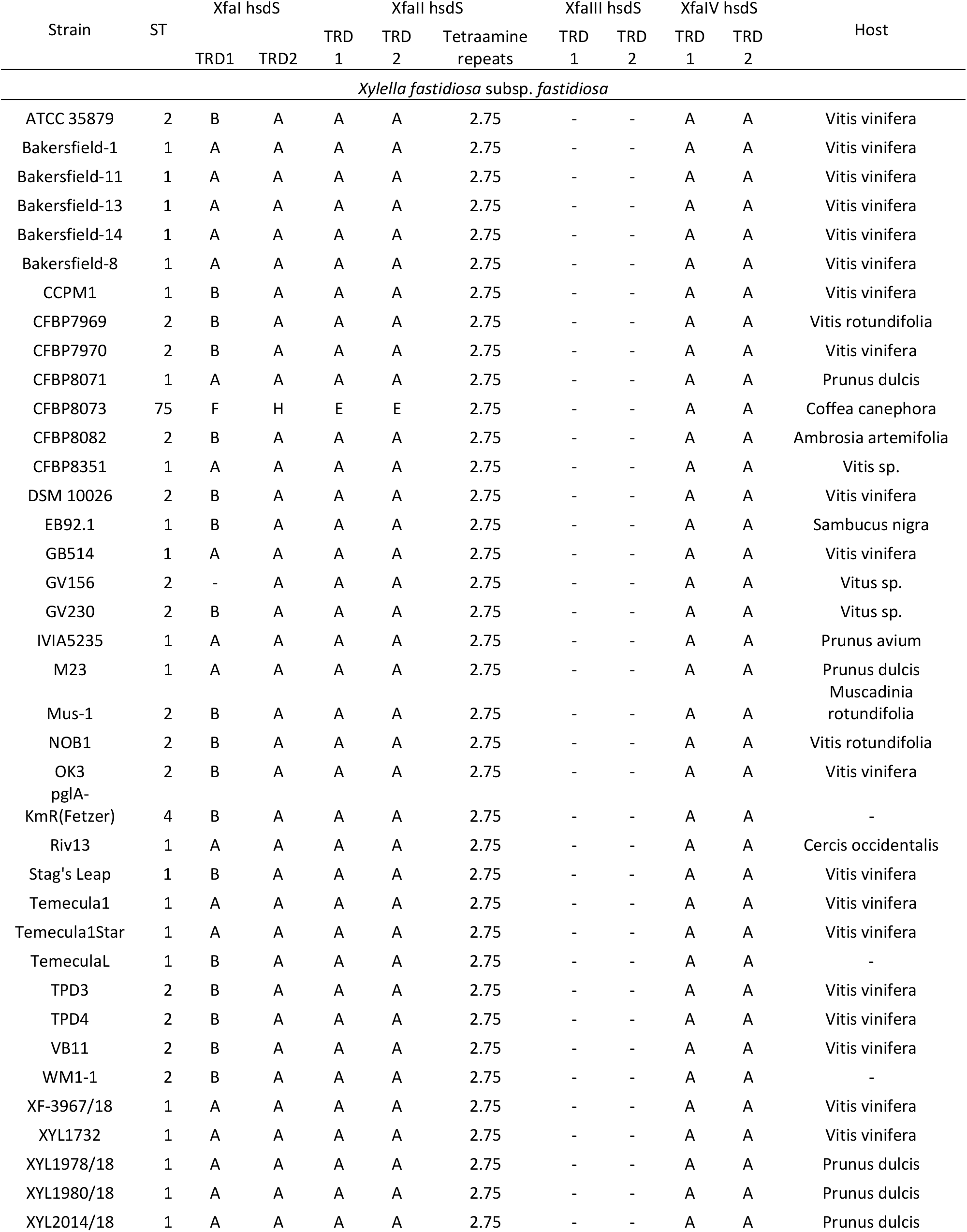
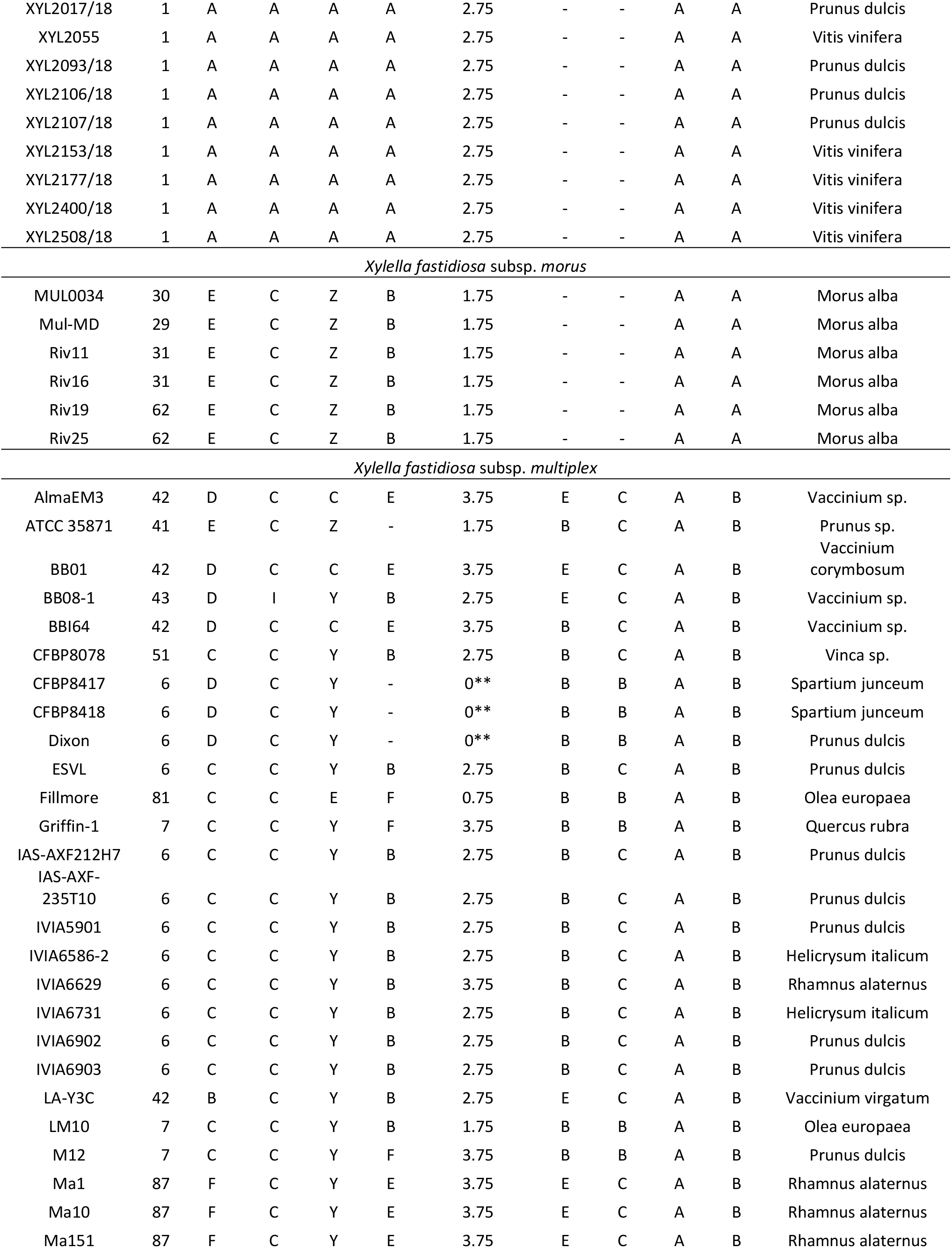
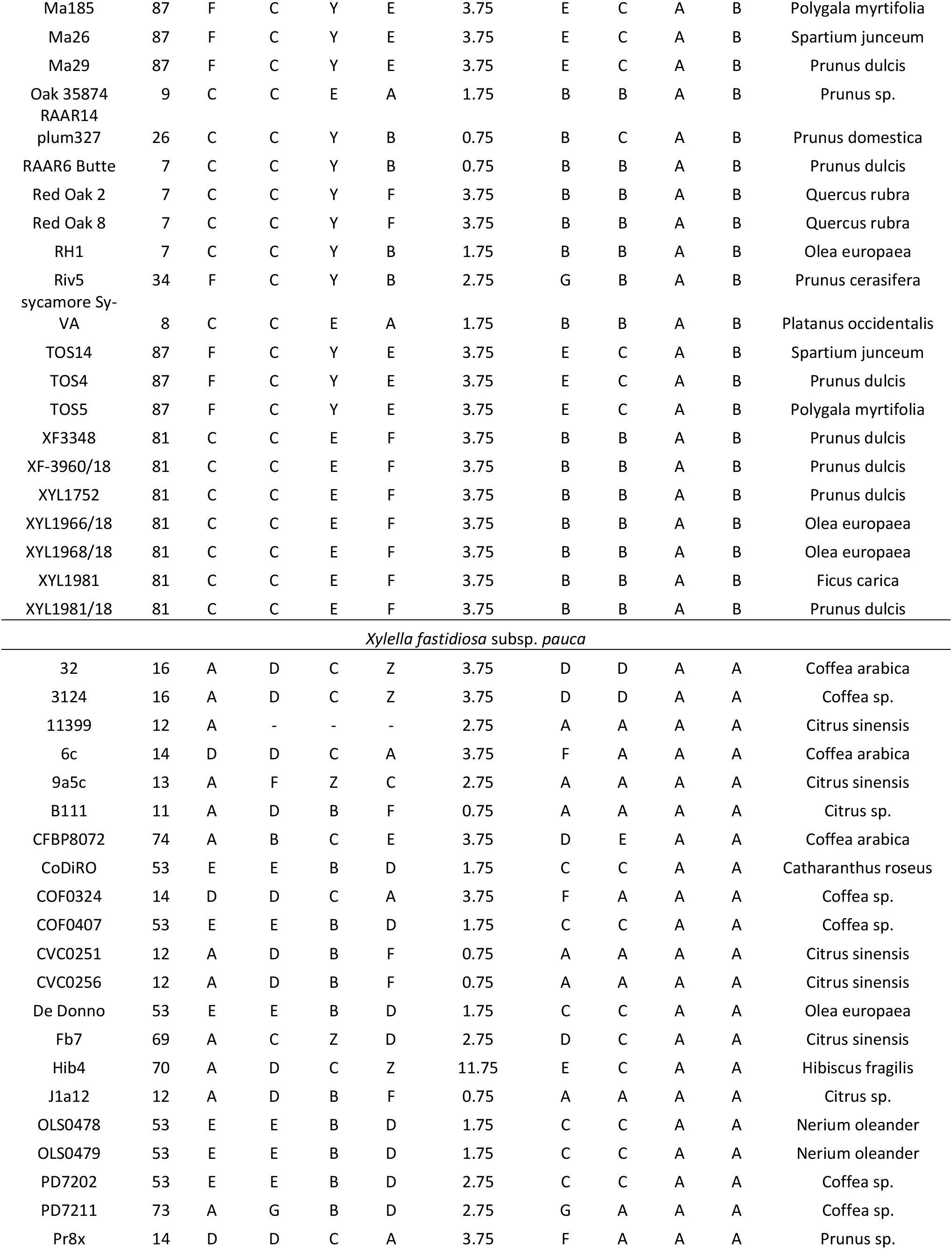
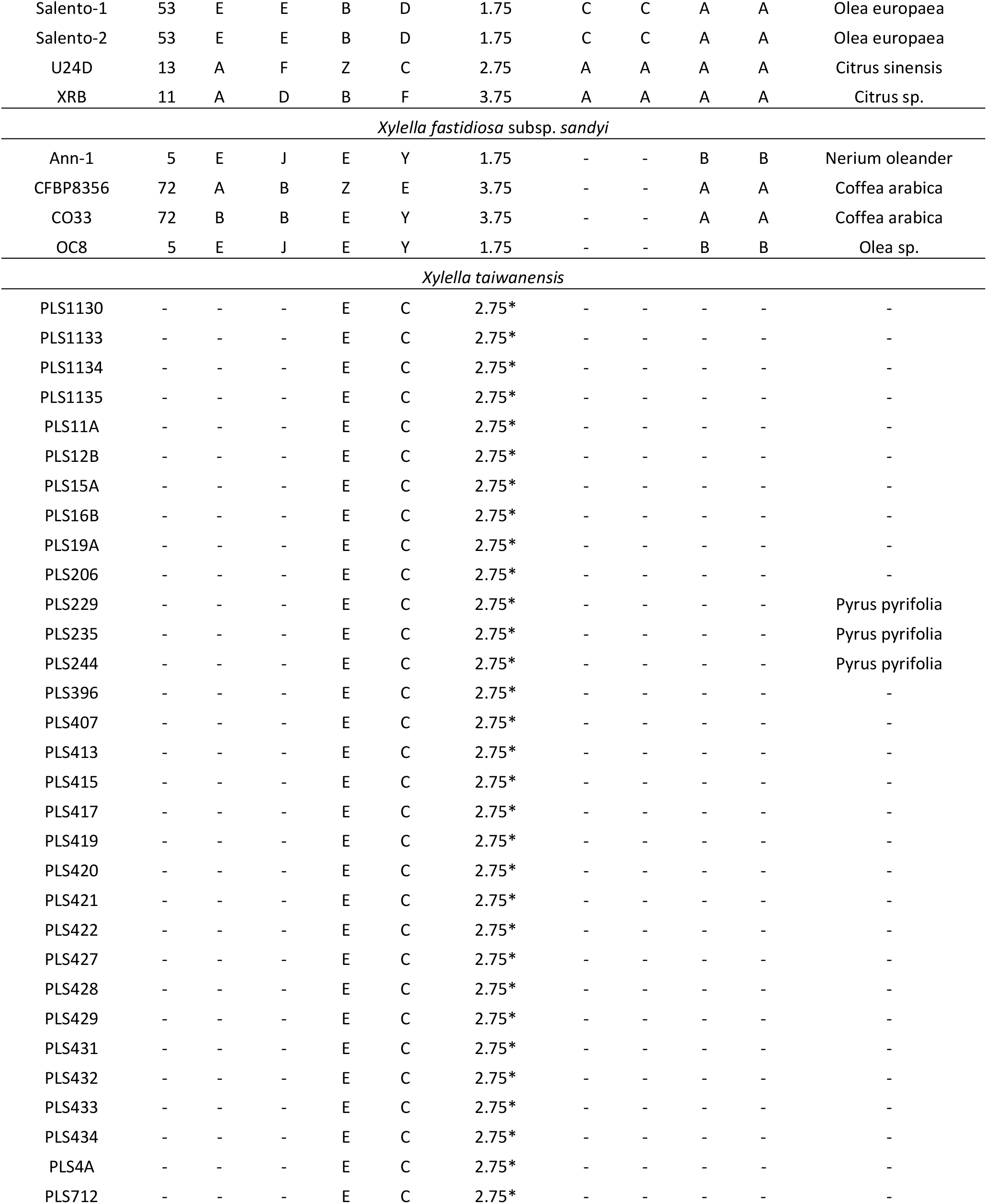
TRD domains identified among *Xylella* strains. ST = multi-locus sequence type. “*” Indicates the core conserved region of these XfaII *hsdS* genes contain an “LHAE” tetraamine repeat instead of “LEAE”. “**” indicates no core conserved region could be identified for XfaII *hsdS* in these strains.

### *HsdS* alleles are variable within and across subspecies

Type I R-M systems can rapidly evolve new methylation and restriction specificities by exchanging TRDs between different *hsdS* alleles (i.e., recombination). We assessed the variation within *hsdS* genes of *X. fastidiosa* by classifying TRD-1 and TRD-2 of each Type I R-M system into homology groups for that specific TRD (e.g., TRD-1 “A”, TRD-1 “B”, etc.). Unique alleles are designated by their TRD-1 and TRD-2 classification (e.g., “AA”, “BA”, etc.), and the combination of these alleles present in a given strain determines its *hsdS* allele profile. Between two and ten variants of each TRD were identified among *hsdS* genes belonging to each Type I R-M system examined in this study (Table 2, Figure 2); in total, 44 unique TRD sequences were identified. No TRD sequences were shared between different R-M systems (e.g., XfaI and XfaII, etc.). In XfaI, XfaIII, and XfaIV, each TRD variant is found only in TRD-1 or TRD-2. In XfaII, two TRDs (“Y” and “Z”) can be found in either TRD-1 or TRD-2, suggesting this domain shuffled positions. TRDs are arranged in combinations yielding 3-18 unique alleles per Type I R-M system, excluding partial or fragmented alleles (Figure 3). In total, 49 unique complete *hsdS* alleles and four incomplete *hsdS* alleles with a single TRD domain were identified associated with XfaI-IV across all *X. fastidiosa* strains, in addition to the single XfaII *hsdS* allele found only in *X. taiwanensis*. Only five alleles were shared between *X. fastidiosa* subspecies. Within XfaI, allele “AB” is shared between *pauca* and *sandyi*, while “EC” is shared between *morus* and *multiplex*. Within XfaII, allele “CE” is shared between *pauca* and *multiplex*. Within XfaIII, allele “EC” is shared between *pauca* and *multiplex* strains. Within XfaIV, allele “AA” is found in all *fastidiosa, morus*, and *pauca* strains, in addition to two *sandyi* strains. Incomplete *hsdS* were identified within XfaI or XfaII in 6/132 strains.

**Figure 2.**
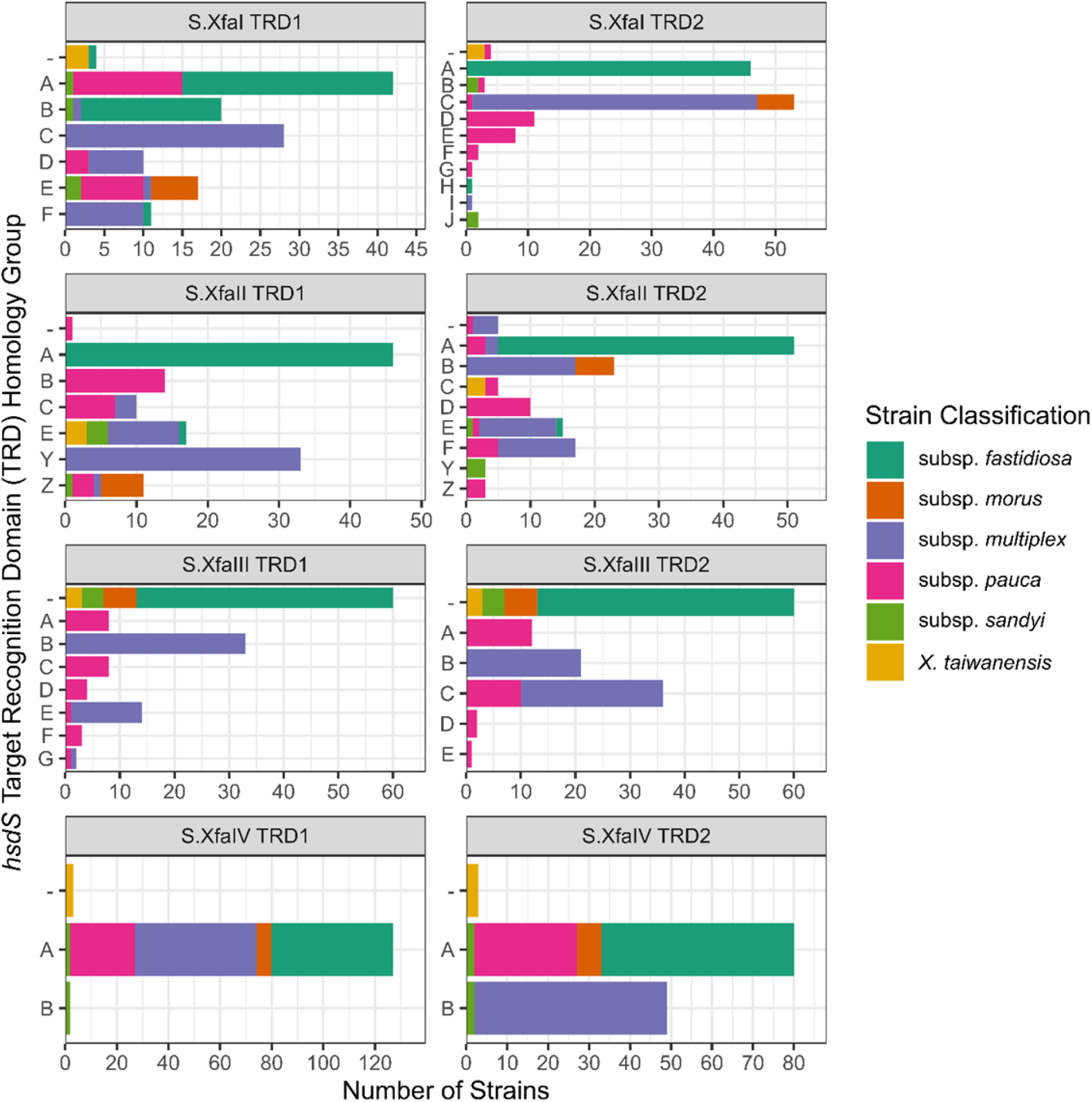
Target recognition domains are shared among *Xylella* strains. Target recognition domain (TRD) abundance across *Xylella fastidiosa* subspecies and *X. taiwanensis* strains. TRDs are presented for each position (e.g., TRD1 and TRD2) in the specificity subunit associated with Type I R-M systems identified in *Xylella fastidiosa* (e.g., S.XfaI, S.XfaII, S.XfaIII, and S.XfaIV). Colors indicate *Xylella fastidiosa* subspecies and *X. taiwanensis*. S.XfaII TRDs “Y” and “Z” are equivalent between TRD-1 and TRD-2; all other TRDs are unique to that TRD position and R-M system. “-” indicates no TRD domain was identified for this system.

**Figure 3.**
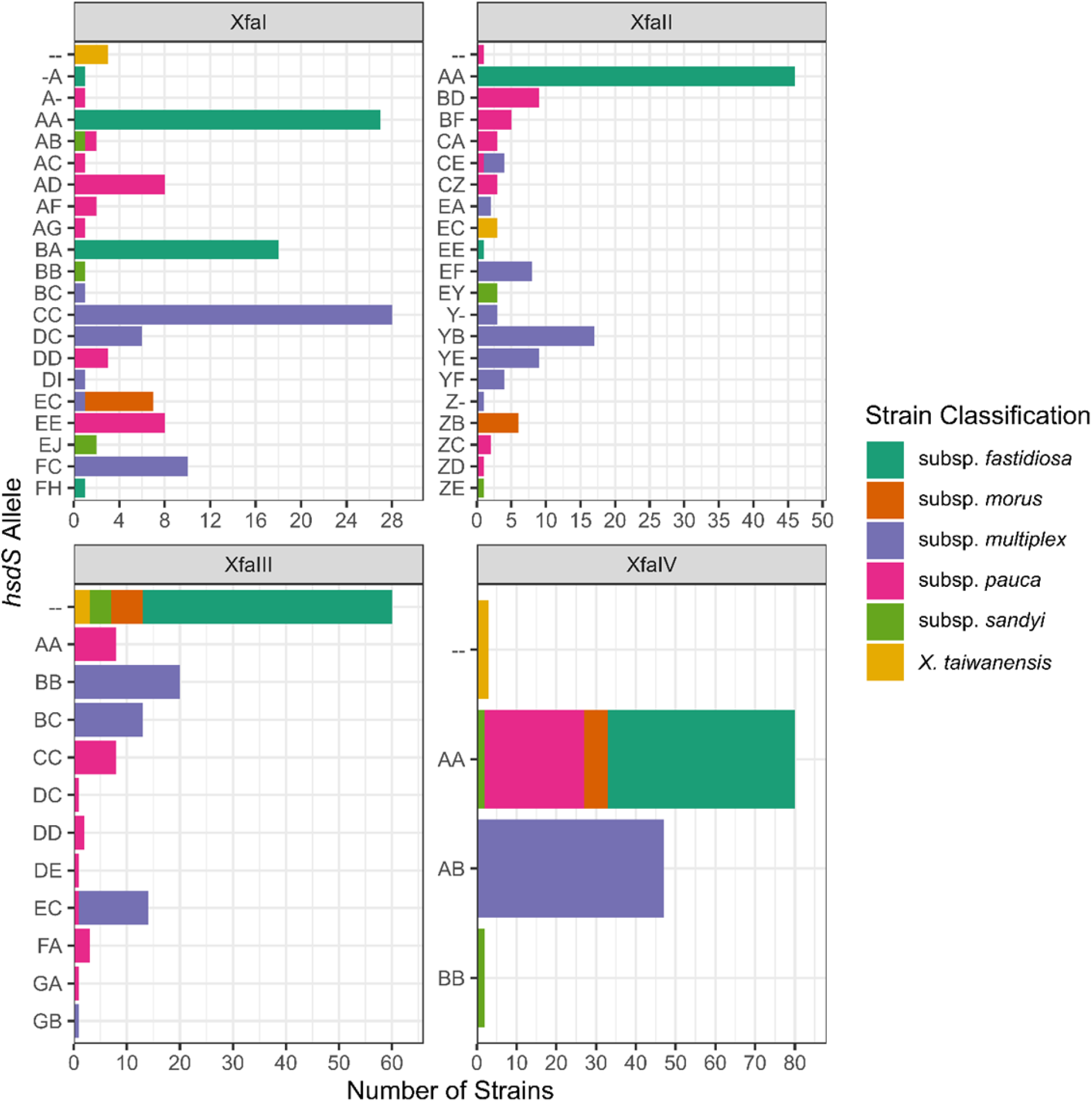
Unique *hsdS* alleles are rarely shared across *Xylella* strains. Type I R-M system specificity subunit (e.g., *hsdS*) alleles identified across *Xylella fastidiosa* subspecies and *X. taiwanensis* strains for XfaI, XfaII, XfaIII, and XfaIV. *HsdS* alleles consist of two target recognition domains (TRDs) and are presented as a combination TRDs (i.e., an allele with TRD1 = B and TRD2 = A is presented as “BA”). Each TRD type is unique to each R-M system (e.g., XfaI “AA” and XfaII “AA” are not equivalent), and with the exception of XfaII TRDs “Y” and “Z”, each other TRD position (e.g., TRD-1 and TRD-2). “-” indicates no TRD was identified in this position.

### Specific *HsdS* allele profiles are predominantly uniform within and limited to monophyletic clades

*HsdS* allele profiles reflect strain relatedness as determined by whole-genome phylogenetic analysis, with strains sharing an allele profile typically forming a monophyletic cluster, or occasionally a paraphyletic cluster (i.e., not including all descendants). In total, 31 combinations of alleles (allele profiles) were identified among *X. fastidiosa* strains, and one among *X. taiwanensis* strains (Figure 4, Figure S1). This variation is most apparent between subspecies, with few TRDs and no alleles shared across subspecies boundaries. Furthermore, each *X. fastidiosa* subspecies, with the exception of subsp. *morus* strains, has multiple *hsdS* allele profiles. Often this is consistent with ST, but not always (Figure 4). Typically, each ST has one to two unique allele profiles, and where allele profiles are shared across more than one ST, both STs and allele profiles form a monophyletic clade (i.e., subsp. *morus* STs, subsp. *pauca* ST 11 and ST 12). A few cases were identified where allele profiles exist across multiple STs, or are paraphyletic with respect to the whole genome phylogeny. A similar pattern of *hsdS* variation between clades has been observed in *Staphylococcus aureus*, where Type I R-M systems restrict gene flow between lineages (34).

**Figure 4.**
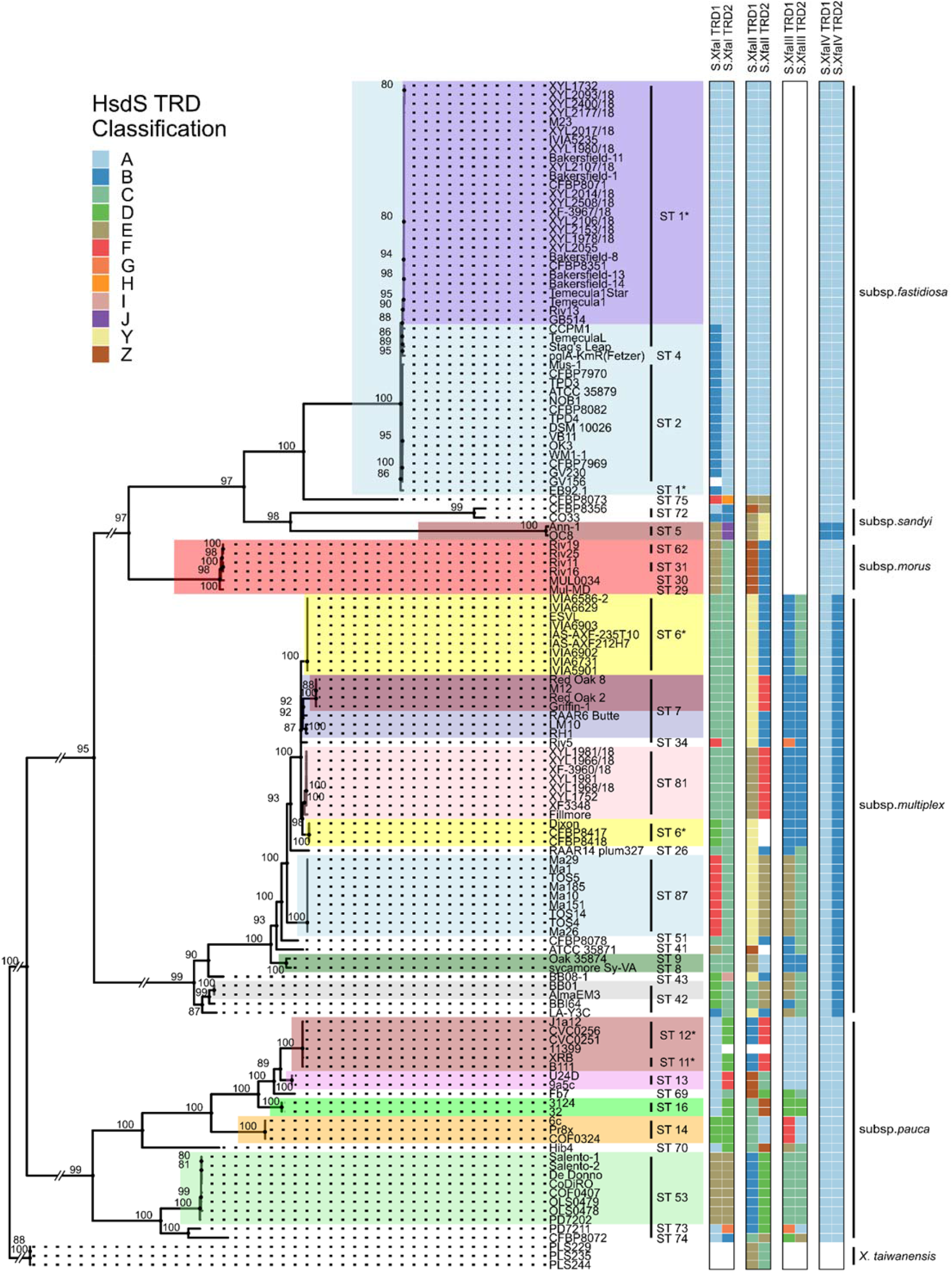
*HsdS* alleles varies among genetic lineages based on combination of target recognition domain sequences TRD1 and TRD2. Long branches leading to X. taiwanensis and X. fastidiosa subspecies were rescaled and are indicated by parallel slashes. Bootstrap values (n=1000) are indicated by each node. *HsdS* alleles are presented as a combination of the constitute target recognition domains (TRDs). TRDs for the single *hsdS* gene in each system are indicated (e.g., S.XfaI TRD1 and S.XfaI TRD2). TRDs were given a letter designation indicating their homology group per domain; letters between R-M systems are not equivalent, nor are letters between TRDs equivalent, with the exception of TRDs “Y” and “Z” in XfaII, which were identified in both TRD1 or TRD2. Sequence type (ST) and subspecies groups are indicated. Clades with a uniform allele type are highlighted.

Among subsp. *fastidiosa* strains, three allele profiles were identified, with three alleles for XfaI and two alleles for XfaII. The majority of subsp. *fastidiosa* strains are distinguished by a difference in TRD-1 of XfaI, having either TRD “A” or “B”. All “AA/AA/--/AA” strains belong to ST1, and form a monophyletic clade containing most California subsp. *fastidiosa* strains as well as strains likely introduced to Europe from California. A paraphyletic group, including all ST2, ST4, and four ST1 strains, have the “BA/AA/--/AA” allele profile. Thus, it is plausible that the “AA/AA/--/AA” profile shared by most ST1 strains arose as the result of XfaI *hsdS* recombination in a common ancestor from the “BA/AA/--/AA” lineage. The single representative of ST 75, isolated in France from a *Coffea canephora* plant imported from Mexico (35), shares no TRDs with the other subsp. *fastidiosa* strains in XfaI or XfaII, but does share one TRD in both XfaI and XfaII with ST87 subsp. *multiplex* strains, and one TRD in XfaII with *sandyi* strains.

Substantially more variation is present among subsp. *pauca hsdS* alleles than in subsp. *fastidiosa*. Ten allele profiles were identified, including 8 XfaI alleles (1 partial), 7 XfaII alleles, and 8 XfaIII alleles. Strains belonging to ST 11 and 12 form a monophyletic clade and share an allele profile, except for strain 11399 which contains fragmented *hsdS* components with no identifiable TRDs for XfaI and XfaII. Each other ST has its own unique *hsdS* profile.

Strains from subsp. *sandyi* have three allele profiles, composed of three XfaI alleles, two XfaII alleles, and two XfaIV alleles. The two ST5 strains, both isolated from plant material in Southern California, have a uniform profile. In contrast, the two ST72 strains have different alleles in XfaI and XfaII, with the XfaI allele of strain CFBP8356 shared with CFBP8072, a subsp. *pauca* ST74 strain.

Strains belonging to subsp. *multiplex* have the most diverse *hsdS* allele profiles, with 6 alleles at XfaI, 8 alleles at XfaII (including 2 partial alleles), and 4 alleles at XfaIII, arranged in 13 allele profiles. While some clearly defined clades (e.g., ST 87) have unique *hsdS* allele profiles, multiple STs in subsp. *multiplex* contain two distinct allele profiles. While ST7 strains are a monophyletic lineage, two sub-lineages are present that differ in TRD2 of XfaII. In contrast, ST6 strains are distributed across two polyphyletic lineages, each with their own unique *hsdS* allele profile that differ in at least one TRD domain of XfaI, XfaII, and XfaIII. Notably, RAAR14 plum327 (ST26) and CFBP8078 (ST51) share the same allele profile (“CC/YB/BC/AB”) as ST 6 strains isolated from European countries, despite being a paraphyletic group isolated from different continents. This paraphyletic distribution suggests this allele profile may have present in the common ancestor of these strains and was altered through recombination in other *X. fastidiosa* subsp. *multiplex* lineages, or that recombination among strains has resulted in this profile being moved between or generated within multiple lineages. An unexpected amount of diversity among *hsdS* subunits was identified among blueberry-infecting ST 42 strains, with three unique allele profiles represented across four strains in a manner consistent with their phylogeny. Strains in this clade are considered to undergo inter-subspecies recombination, which may explain the diversity in these systems. Given the substantial *hsdS* variation within these strains and subsp. *multiplex* generally, it is possible that *hsdS* allele variation provides an adaptative advantage to new or changing environments (36).

### Geographic origin is not a sole predictor of *hsdS* allele profile

There are several examples of *X. fastidiosa* subsp. *fastidiosa*, subsp. *multiplex*, or subsp. *morus* strains with different geographic origins (e.g., United States vs. Europe) sharing the same allele profile (Table 2, Table S3). However, *X. fastidiosa* strains that were collected from the same geographic location do not necessarily share the same allele profile (e.g., subsp. *fastidiosa* strain M23 and subsp. *multiplex* strain M12). Furthermore, some allele profiles are remarkably consistent within a geographic region, such as those of *X. fastidiosa* subsp. *pauca* ST53 responsible for olive decline in Italy, *X. fastidiosa* subsp. *fastidiosa* ST1 in the California San Joaquin Valley, *X. fastidiosa* subsp. *multiplex* ST87 in Tuscany, Italy, or *X. fastidiosa* subsp. *multiplex* ST81 in the Balearic Islands in Spain. The majority of strains examined in this study are associated with disease outbreaks in specific geographic regions as the result of an introduction event, which can cause a genetic bottleneck and limit diversity. Although it is possible this trend is due to sampling bias, it is reasonable to assume that the lack of diversity among Type I R-M system *hsdS* components in many geographic regions is due to a combination of 1) limited diversity within introduced strains and 2) few or no locally available compatible strains to donate novel *hsdS* alleles. It is possible that greater diversity among *hsdS* alleles would be observed within monophyletic clades if multiple *X. fastidiosa* strains from distinct lineages were present in the same geographic region, or if strains from *X. fastidiosa’s* native range or a wider variety of hosts were sampled.

### Detectable methylation patterns are consistent with *hsdS* allele profiles, not sequence type or subspecies

Genetic variation among *X. fastidiosa hsdS* subunits suggests strains differentially modify specific DNA motifs, which could result in distinct epigenetic profiles across strains or subspecies. Nanopore sequencing was used to compare DNA methylation motifs of 20 wild-type *X. fastidiosa* strains representing 13 allele profiles (Fig. 3). Modified DNA motifs were detected from Nanopore reads using Nanodisco (37). One to six distinct modified DNA motifs were identified per strain, up to three of which are associated with Type I R-M system activity. Bipartite motifs were typically predicted to have a 6mA modification, whereas other motifs had either 6mA, 5mC, or 4mC modifications. In most strains, the reverse complement of non-palindromic motifs were identified. Nanodisco does not determine motif directionality, (i.e., 5’ to 3’ vs. 3’ to 5’), thus these were assumed to be different orientations of the same motif and a direction was arbitrarily chosen. As expected, the specific Type I R-M system associated motifs identified in each strain are consistent with variation among *hsdS* alleles encoded in that strain, instead of sequence type or subspecies. In total, 12 combinations of Type I R-M system modifications were identified from 14 allele profiles. In two strains, OC8 and Riv19, no Type I R-M system associated motifs were detected. Many of the detected modification motifs were shared between strains of the same subspecies and corresponded to shared *hsdS* alleles. Likewise, methylation motifs from Type I R-M systems were not necessarily consistent with STs, but were consistent within an *hsdS* allele profile.

Only in one case did strains with different allele profiles share the same motif pattern: *X. fastidiosa* subsp. *multiplex* strains LM10, RH1, and Fillmore. These strains share the same *hsdS* alleles at all sites except XfaII, and only two Type I R-M associated motifs were identified; thus, the most plausible explanation is that XfaII is not active in these strains. Furthermore, although LM10 and RH1 share their XfaII *hsdS* allele (“YB”) with Riv5, Y3C, and BB08-1, none of the bipartite motifs detected in LM10 and RH1 correspond to motifs detected in Riv5, Y3C, or BB08-1. Although LM10, RH1, and Filmore were recovered from the same host (olive) and from the same greater geographical region (Southern California, USA), many more strains would need to be examined to determine if independent loss of function in XfaII is common to subsp. *multiplex* strains recovered from olive or southern California, or simply coincidental among these strains.

The majority of subsp. *fastidiosa* strains have one of two allele profiles, which differ from each other only in TRD1 of XfaI, represented among this set of strains by Stag’s Leap (“BA/AA/--/AA”) and the remaining ST 1 subsp. *fastidiosa* strains (“AA/AA/--/AA”) (Figure 4). Because these strains have identical allele profiles except for a single TRD substitution, methylated motif analysis should identify one bipartite motif where Stag’s Leap and the other ST1 strains share sequence in only one of the two halves, and one motif matching this criteria was identified (“G6mATT(N_7_)TCC” in Stag’s Leap and “G6mATG(N_7_)TCC” in other ST 1 *fastidiosa* strains). Deletion mutants of XfaI *hsdS* genes were generated in Stag’s Leap (LZ752_RS11070) and ST1 strain M23 (XFASM23_RS10960) to verify that these differential motifs are associated with XfaI in these strains. Analysis of both Stag’s Leap and M23 XfaI *hsdS* mutants with Nanodisco failed to identify these differential motifs. Thus, the methylated motifs “G6mATG(N_7_)TCC” and “G6mATT(N_7_)TCC” can be assigned to XfaI alleles “AA” and “BA”, respectively.

Seven additional motifs associated with either Type II or Type III R-M systems were identified as well (Figure 5). Unlike Type I R-M system associated motifs, these motifs are substantially less diverse and several are commonly shared among members of a subspecies. One such motif is “TTCGA6mA”, the expected recognition site of a Type II R-M system (PD1667-PD1668) homologous to NspV present in some *X. fastidiosa* subsp. *fastidiosa* strains (16). Although this system was previously described as specific to subsp. *fastidiosa*, both PD1667-PD1668 and the “TTCGA6mA” motif were identified in subsp. *multiplex* strain BB08-1 and all three subsp. *morus* strains. Strikingly, the motif “CGTA4mCG” was identified in all subsp. *multiplex* strains included in this analysis. One motif likely belonging to a Type III R-M system was identified in both subsp. *fastidiosa* strains and the sole evaluated subsp. *pauca* strain, De Donno. A discrepancy we observed among these additional motifs was the detection of the motif “AT5mCGAT” in the M23ΔXfaIIHsdS mutant, despite this motif not being detected in wild-type M23. One explanation could be that this palindromic motif overlaps with the first three bases of the target motif of the XfaII *hsdS* allele in this strain, “G6mATG(N_7_)TCC”, and is thus is not detected in most samples by the analysis method we used.

**Figure 5.**
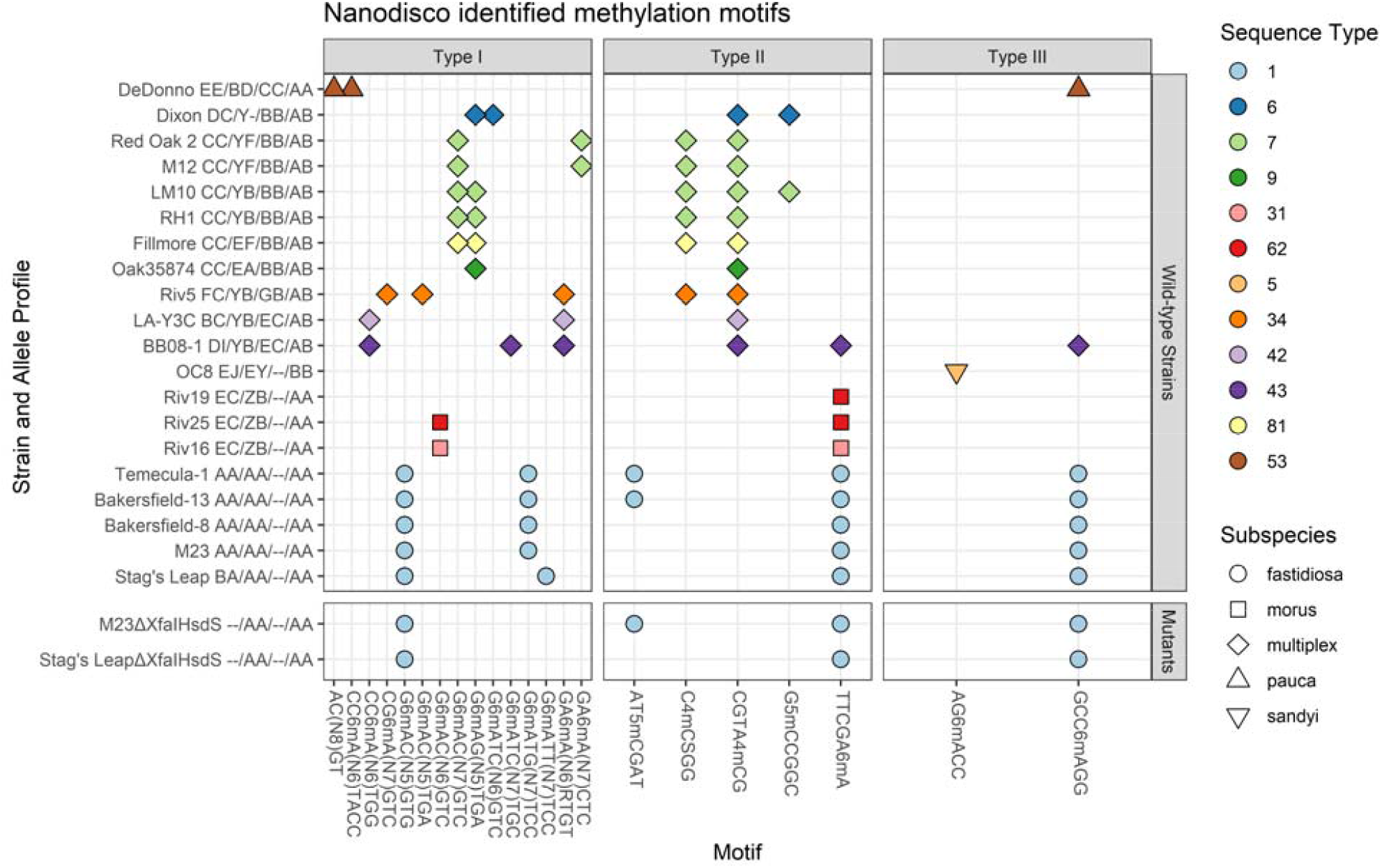
Methylated motifs detected in *X. fastidiosa* strains. Nanodisco-identified methylated motifs detected in *X. fastidiosa* strains. Type I R-M system *hsdS* allele profile is indicated after each strain name. Sequence type (ST) and subspecies are indicated by color and shape, respectively. Detected motifs are presented along the x-axis, separated into categories by the expected class of system responsible for the motif (e.g., Type I-III). Type I R-M system spacers are indicated in parenthesis as “Nn”, where n indicates the length of the spacer (e.g., N_5_ = NNNNN).

### Type I R-M systems in *Xylella* strains frequently have disruptive mutations

In all strains evaluated for methylation patterns, fewer modified bipartite motifs were detected than Type I R-M systems encoded in each genome, suggesting some Type I R-M systems may not be functional in individual strains. Type I R-M systems can be inactivated by mutations which disrupt *hsdS* or *hsdM*, preventing both restriction and methylation activity, while disruptive mutations in *hsdR* prevent restriction activity but have no effect on methylation. We manually examined Type I R-M system *hsdS, hsdM*, and *hsdR* components in each of the 20 strains evaluated for methylation to determine if coding sequences of these components from each system were intact or disrupted, and identified disruptive mutations in one or more Type I R-M system components in several strains (Figure 1; Figure 6). These disruptive mutations typically introduce a frameshift resulting in mistranslation or a premature stop codon (e.g., XfaIII-*hsdM* in M12, XfaI-*hsdS* and XfaII-*hsdS* in OC8) or in some cases are large deletions (e.g., loss of XfaII-*hsdM* in *X. fastidiosa* subsp. *morus* strains), suggesting these Type I R-M systems are likely not functional for methylation and/or restriction in those strains. Among the 20 strains examined, the least putative functional variation was observed among XfaI and XfaIV systems. All XfaI components are intact in 18/20 strains, while only 1/20 strains had any intact XfaIV components (Figure 6). Among strains carrying XfaIII, 5/11 had disrupted *hsdM* (2/11) or *hsdR* (3/11) components, suggesting XfaIII may be methylating DNA in the absence of a functional restriction component in some cases. The most strain-to-strain variation among Type I R-M system components was observed in XfaII, which has a disruptive mutation in *hsdS* coding sequences in 9/20 strains examined. It is worth noting *X. fastidiosa* subsp. *multiplex* strains included in this analysis have the most strain-to-strain variation in Type I R-M system component integrity across all systems, particularly in contrast to subsp. *fastidiosa* and *morus* where no variation was observed. Furthermore, subsp. *multiplex* ST7 strains (i.e., Red Oak 2, M12, LM10, and RH1) have three distinct mutation patterns expected to result in differential activity of XfaII and XfaIII. However, although the low number of representatives from each ST and subspecies makes it unclear how widespread or consistently inactivating mutations appear within a ST or subspecies.

**Figure 6.**
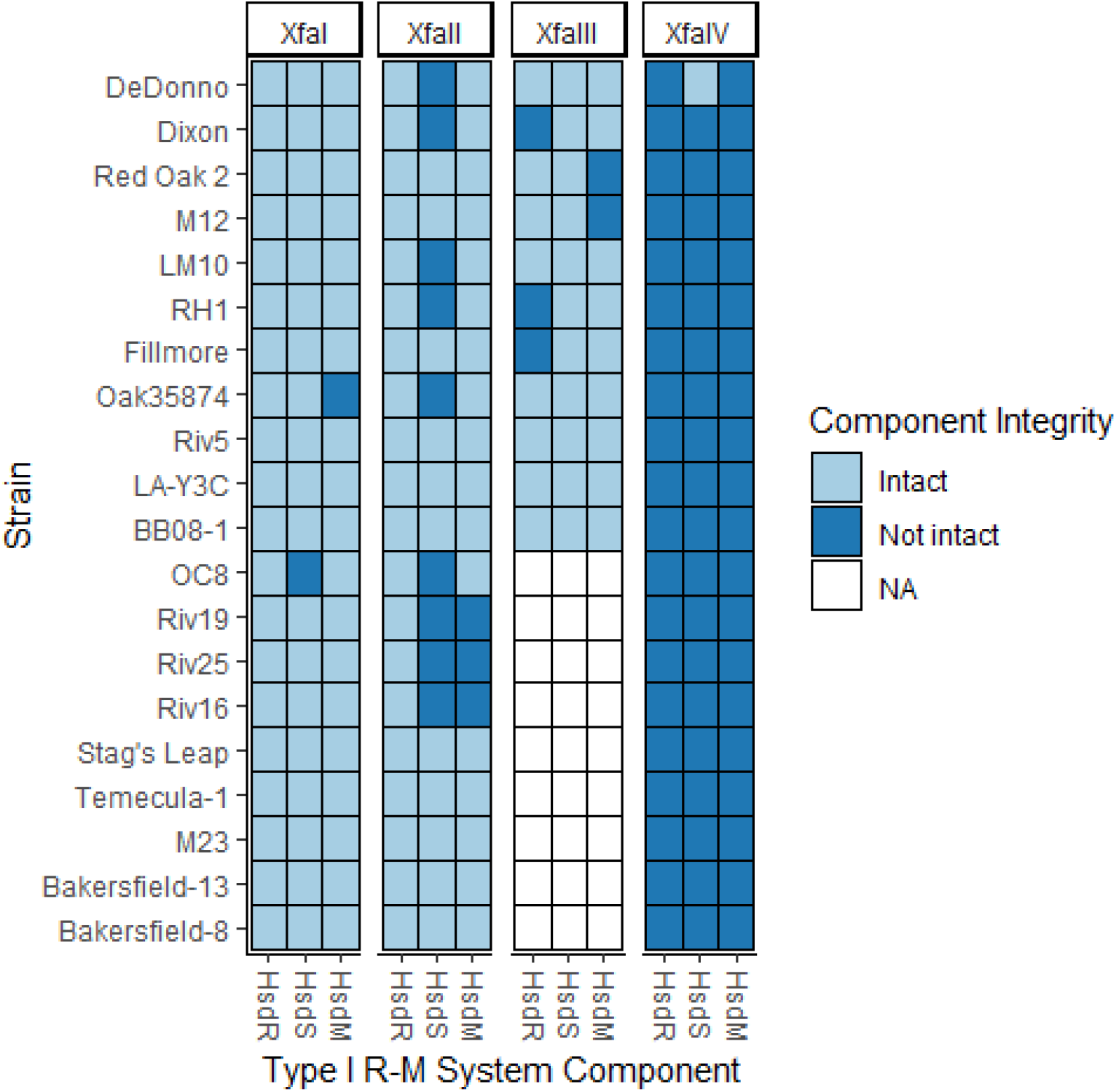
Type I R-M system components are frequently disrupted in *X. fastidiosa* strains. Type I R-M systems require *hsdS* and *hsdM* genes for both restriction and modification; systems without a functional *hsdR* component can still modify DNA. Intact = full length coding sequence present; Not Intact = component coding sequence disrupted or truncated; NA = system not present.

Out of the 20 strains examined, one strain of *X. fastidiosa* subsp. *morus* (Riv19) and two subsp. *multiplex* strains (LA-Y3C and Fillmore) had a discrepancy between number of intact (e.g., putatively functional) Type I R-M systems and number of bipartite motifs identified (Figure 5, Figure 6). It is not clear if this is due to differences in gene expression or a limitation of our detection method. Overall, Type I R-M systems in *X. fastidiosa* often appear to be inactivated. This could contribute to *X. fastidiosa* natural competence, as Type I restriction inhibitors are known to improve transformation efficiency in some strains (16, 38). Inactivation of different Type I R-M systems could also contribute to the variability in natural competence observed between *X. fastidiosa* strains and subspecies (14).

Combining intact Type I R-M systems, allele profiles, and detected bipartite motif results across strains allows some Type I R-M system modification motifs to be tentatively assigned to specific alleles by process of elimination. For example, GA6mA(N_6_)RTGT is likely the target motif of XfaII-*hsdS* allele “YB”, because this motif is present in all strains with an intact XfaII system and “YB” allele. Similarly, the motif G6mAC(N_5_)TGA was detected in all strains with a functional XfaIII system with allele “BB”, and thus is likely the target of XfaII allele “BB”. However, even tentatively assigning a target motif to every allele remains challenging due to the variation among these systems, and inability to determine directionality of a motif and thus assign each half of a bipartite recognition motif to specific TRDs. Definitively assigning motifs to specific Type I R-M systems and alleles requires additional experimental evidence (e.g., knockout mutants) and/or genomic methylation analysis of strains with single TRD differences across an otherwise conserved allele profile, such as with XfaI-*hsdS* alleles “AA” and “BA”.

### XfaII *hsdS* subunits contain multiple SSR tracts with variable lengths suggesting potential for phase variation

Phase variation, the dichotomous switching of bacterial phenotypes between different states, has been implicated in a wide range of bacterial adaptation strategies including those important for host colonization (39). In several animal pathogenic bacteria, phase-variable restriction-modification systems have been identified that cause changes in methylation state in across the genome, which in turn affects the expression of multiple genes, including those involved in environmental adaptation or virulence (40). Little information exists regarding phase variation in *X. fastidiosa*, though it has been speculated to be responsible low frequency changes in colony morphology after repeated subculturing (41). However, dramatic phenotypic shifts are necessary in the lifecycle of *X. fastidiosa* as it transitions between the plant host and insect vectors (42), and it is possible that phase variation contributes to this at some stage. Type I R-M systems can principally exhibit phase-variation through either recombination between multiple *hsdS* genes within a single R-M system, or on/off switching of *hsdM* or *hsdS* subunits mediated by SSR length variation. We did not identify multiple *hsdS* genes associated with any Type I R-M system among *X. fastidiosa* strains included in this study, thus *hsdS* recombination-mediated phase variation unlikely to occur (40). However, three variable-length SSRs were consistently identified in XfaII *hsdS* across strains (Figure S2). One SSR, a repeating sequence of twelve bases, is located within the core conserved region of XfaII *hsdS*, resulting in a tandem tetra-amino-acid repeat (“LEAE”) in the core conserved region of the HsdS protein subunit. The length of the LEAE repeat motif varies between strains, the “LEAE” motif varies between strains typically has 0.75-3.75 repeats (e.g., “LEA”, “LEAELEA”, “LEAELEAELEA”, etc.), with subsp. *pauca* strain Hib4 encoding an unusual 11.75 repeats (Table 2). “LEAE” repeat length not only varies between but within XfaII *hsdS* alleles, notably allele “YB”. Strains with this allele may contain 0.75 (RAAR14 Plum), 1.75 (LM10 and RH1), 2.75 (Riv5 and IVIA5901), or 3.75 (IVIA6629) LEAE repeats. Tetraamine repeats within the core conserved region are a common feature of the Type I R-M system family IC *hsdS* subunits; the length of the repeat influences the length of the bipartite DNA recognition motif nonspecific spacer and, in some cases, may alter the DNA recognition sequence of TRD2, while the specific sequence of this repeat motif serves to separate such *hsdS* subunits into compatibility groups within the type IC family (20, 43). The significance of these alleles containing less than a full repeat in *X. fastidiosa* is not clear, however mutants that entirely lack tetraamine repeats in a similar Type I R-M system in *Neisseria gonorrhoeae* lose R-M activity associated with that system (43). This may explain the discrepancy between the number of detected bipartite motifs and intact Type I R-M systems in strain Fillmore, which carries an XfaII “EF” allele (QJP49250.1) with 0.75 “LEAE” repeats. Other strains with this allele have three full LEAE repeats, although their methylated motifs are not known. *X. taiwanensis* strains have a similar tetraamine repeat motif in XfaII *hsdS*, translating to “LHAE” with an invariable 2.75 repeats across 31 strains, suggesting that from *X. taiwanensis* XfaII *hsdS* may not be compatible with XfaII systems from *X. fastidiosa*. It is not clear why XfaII *hsdS* tetraamine are a uniform length within *X. taiwanensis* while varying in *X. fastidiosa*, and could be indictive of differing selective pressures on these species or simply the greater genetic diversity present in available *X. fastidiosa* genomes.

The remaining two SSRs in XfaII *hsdS* are both poly-cytosine SSR tracts containing 7-9 C’s. One tract immediately precedes the core conserved region in strains encoding alleles where TRD1 = “E”, “Y”, or “Z”. Among subsp. *multiplex* ST7 strains, in-frame versions of alleles containing this tract have 8 C’s (i.e., in RAAR6 Butte, M12, Red Oak 2, Red Oak 8, and Griffin-1), while loss of one C within this tract introduces frameshift mutations in ST7 strains LM10 and RH1. Similarly, strain RAAR14 Plum (ST26) has 9 C’s in this tract, resulting in a frameshift mutation in XfaII *hsdS*. The second tract occurs at the beginning of the terminal conserved region in all alleles, and length variation introduces frameshift mutations resulting in mistranslation of the terminal conserved region in strains De Donno and Oak35874. *X. taiwanensis* strains have similar tracts in corresponding locations of XfaII *hsdS*, although both tracts are split by a substitution mutation (C to A or T, respectively) and did not vary in length across strains.

Although SSRs within XfaII *hsdS* exhibit variable lengths across strains, it remains to be determined if XfaII is a phase-variable Type I R-M system. Strains with SSR-associated frameshifts in XfaII *hsdS* had correspondingly fewer methylated bipartite motifs detected by Nanodisco (e.g., De Donno, LM10, RH1, and Oak35874), suggesting changes in these tracts may allow some strains to modulation of XfaII R-M activity. However, it is not clear if changes in SSR length occur frequently enough in *X. fastidiosa* to create distinct lineages within a population existing within a host or in culture. It is plausible that these systems are reversibly disrupted infrequently as an adaptive characteristic on an evolutionary time scale, or perhaps as a response to environmental shifts (i.e. host colonization, prolonged culturing).

## Conclusion

R-M systems are a common feature in bacteria which can provide a barrier to horizontal gene transfer and recombination, thus affecting strain evolution. Epigenetic modification can also impact gene expression, and there is growing evidence for the importance of R-M systems in bacterial virulence. Although R-M systems were previously identified as important for transformation efficiency in *X. fastidiosa*, and variation in R-M systems present in different strains has been observed (13, 14, 17), the comprehensive functions of epigenetic modification in this pathogen are not well understood. Here, we showed that *X. fastidiosa* strains carry up to four Type I R-M systems, three of which are located in a cluster. These Type I R-M systems have highly diverse *hsdS* gene alleles that vary between distinct clades of *X. fastidiosa*, but are frequently conserved within monophyletic lineages. We showed that a subset of strains with different allele profiles have differentially methylated DNA motifs, with more closely related strains having similar epigenomes. Although these R-M systems are conserved across every strain, individual components are often not intact, suggesting they are not uniformly functional across all *X. fastidiosa* strains. The abundance of R-M systems would seem at odds with the naturally competent nature of *X. fastidiosa*, as previous studies found both naturally occurring and in vitro recombination between strains as well as numerous prophage integrations (11, 14, 17, 28). However, genetic exchanges can be more common between strains with related R-M systems, as modifications on donor DNA block degradation by restriction endonucleases in recipient strains (19). Similarly, population-level R-M system repertoire variation is useful for limiting the spread of a bacteriophage or other mobile genetic element within that population. Successful infections will result in bacteriophage/mobile elements with the host cell’s epigenic modifications, which are not necessarily compatible with other members of the population (24).

In addition to direct impacts on horizontal gene transfer, epigenetic modifications made by DNA methyltransferases can also influence gene expression. The role of DNA methylation has been studied extensively among certain animal-pathogenic bacteria. In particular, *Streptococcus pneumoniae* has a conserved phase-variable Type I R-M system with multiple *hsdS* alleles that can affect nutrient acquisition, stress tolerance, and virulence (40, 44). In contrast, a single-*hsdS* Type I R-M system deletion mutant in *Streptococcus pyogenes* doesn’t have altered transcription (45), suggesting a regulatory role is not a universal characteristic of Type I R-M systems. However, DNA methylation is poorly studied among plant pathogenic bacteria and limited examples of methylation affecting gene expression exist. Altered methyltransferase expression can affect virulence and growth in *Xanthomonas axonopodis* pv. *glycines* and *Xanthomonas euvesicatoria*, although in neither case are these methyltransferases associated with a Type I R-M system (46, 47). In *X. fastidiosa* further work is necessary to understand the functional role of Type I R-M systems both in transcriptional regulation and in plant disease development or environmental adaptation. This characterization of R-M systems and methylation patterns across a wide range of *X. fastidiosa* strains will facilitate future studies investigating the role of epigenetic modification in biology and evolution of this important pathogen. Importantly, further research on epigenetic modification could potentially lead to a better understanding of *X. fastidiosa* phenotypic variation that in some cases is still very poorly understood (i.e. host range interactions).

## Materials and Methods

### DNA preparation, sequencing, and genome assembly

*X. fastidiosa* strains were cultured on PD3 agar medium (48) and incubated at 28°C for 5 days for subsp. *fastidiosa* strains, and 7 days for subsp. *multiplex, morus, sandyi*, and *pauca* strains. Plates were scraped and DNA extracted using the Qiagen DNeasy Blood and Tissue kit including a treatment step with 4 µl RNaseA (100 mg / ml). DNA was quantified via Qubit, used to prepare a sequencing library with an Oxford Nanopore ligation sequencing kit (LSK-109, Oxford Nanopore) with native barcoding and expansion kits (NBD-104 and NBD-114) with short fragment buffer (SFB), and sequencing on a MinIon sequencer with an R9.4.1 flow cell. Resulting reads were basecalled with guppy v3.0.3-4.0.15 with parameters *--qscore_filtering --min_qscore 7* using the high accuracy model (Table S4). Short-read libraries were prepared and sequenced either by the University of California, Davis DNA Technologies core facility using the Illumina Novoseq platform with 150 bp paired end reads and an insert size of 500 bp, or by CDGenomics using the Illumina HiSeq 2500 platform with 150 bp paired end reads. Genomes were assembled using nanopore reads with Flye v.2.7.1-v2.9 with parameters –*nano-raw* –*genome-size* 2.5M –*plasmids*, except for OC8 (–*nano-raw* – *genome-size* 2.7M), as well as Stag’s Leap and Riv11 (–*nano-raw* –*genome-size* 2.5M) (49). Flye assemblies were polished with pilon v1.23 up to five times with random subsamples of Illumina reads with estimated 100x coverage generated with Seqtk v1.3, or until no further changes were made, and rotated so that the first position of each genome corresponds to the first base of *dnaA* (50, 51). For Bakersfield-14, Riv13, and Stag’s Leap, input DNA was fragmented to approximately 8 kb with a Covaris g-TUBE prior library preparation. For Stag’s Leap ΔXfaI*hsdS*, barcoding kits were not used. The updated Bakersfield-1 genome was generated with Unicycler v0.4.8-beta (52). For OC8, reads were filtered to 15 kb minimum length with Filtlong v0.2.0 prior to assembly (53). For Stag’s Leap, MiSeq reads were processed with fastp using parameters ‘-q 20, -e 30 --cut_mean_quality 30 --cut_front --cut_tail’, and reads were not subsampled. Assembly quality was evaluated with CheckM, IDEEL, and fastANI was used to determine ANI to *X. fastidiosa* type strain ATCC 35871 (54–56).

### Whole genome phylogeny

Genome sequences for *X. fastidiosa* and *X. taiwanensis* were retrieved from RefSeq. A core genome alignment was generated with Parsnp v1.5.2, converted to multifasta format with HarvestTools (57), and used to generate a recombination-sensitive whole genome phylogeny with Gubbins v2.4.1, using RaxML v 8.2.12 to construct a phylogeny with ‘–GTRCAT’ and evaluate the phylogeny with 1000 bootstrap replicates (58, 59). *X. taiwanensis* strains were used as an outgroup. The resulting phylogeny and genotype data were visualized in R v4.0.3 using the ‘ggtree’ v2.2.4 package (60) and edited with Inkscape.

### Bioinformatics analysis

*HsdS* alleles were identified with blastn analysis using a set of type alleles as a query (S. File 1) against *Xylella* CDS sequences retrieved from RefSeq; in cases where either a partial *hsdS* gene or no *hsdS* gene was identified, the corresponding genome assembly was queried instead. *HsdM* and *hsdR* components were identified by blastn (1e^−25^ cutoff) using *hsdM* and *hsdR* CDS from the Riv5 RefSeq assembly for XfaI-III, and the REBASE predicted nucleotide sequence for XfaIV components. Riv5 was selected because it does not have disrupted CDS belonging to XfaI-III. Gene alignments were made with Clinker (61) and modified in Inkscape. Genome alignments were made and LCBs identified with progressiveMauve (62), using M12 as a reference strain. Doron defense systems were identified with PADLOC (63). Genomes were re-annotated with prokka v1.14.6 (64) prior to Clinker and Mauve analysis. Frameshifts, SSRs, and “LEAE”/”LHAE” repeats were idenfitied by manual examination of *hsdS* sequences. MLST profiles were determined using the *X. fastidiosa* PubMLST database (65).

### TRD identification and *hsdS* allele classification

An initial set of *hsdS* alleles were extracted from the annotation files available in RefSeq for complete *Xylella* genomes. Additional alleles were identified in the remaining genomes by blastn search against a database constructed from nucleotide sequence of predicted gene products using annotation available in RefSeq, and the highest-scoring hit for each combination of R-M system and genome was extracted. CDS were then clustered using CD-HIT with parameters ‘sequence identity cut-off 0.9 -aL 0.9’, and representatives from each clusters compared. For strains in clusters containing truncated *hsdS* genes, or where one or more TRD domains could not be identified, the genome assembly was searched with blastn using cluster representatives and the results manually examined. Representative *hsdS* alleles for each Type I R-M system are available as supplemental files 1-4.

### DNA modification analysis

For each strain, total genomic DNA was amplified using an illustra *GenomiPhi* V2 DNA *Amplification Kit (*Cytiva), purified using AMPure XP beads (Beckman Coulter), and used for library preparation and sequencing as described above. Methylation predictions were done using Nanodisco v.1.0.1, comparing nanopore reads from genome assemblies (‘native’ DNA) and amplified DNA (37). Nanopore reads used for genome assembly and reads from amplified DNA were re-basecalled with guppy v4.0.15 using the high accuracy model. To retrieve single fast5s from multifast5 files from multiplexed sequencing runs the ont-fast5-api scripts fast5_subset and multi_to_single_fast5 (66) were used to search for read names from corresponding fastqs used for genome assembly of each strain (e.g., demultiplexed reads).

### Construction of deletion mutants

*HsdS* deletion mutants of *X. fastidiosa* subsp. *fastidiosa* strains M23 and Stag’s leap were created by homologous recombination as previously described for *X. fastidiosa* (67). Briefly, 1 kb regions directly upstream and downstream of the *hsdS* open reading frame were amplified from genomic DNA of the respective strains and annealed to either side of a chloramphenicol resistance cassette (Tn5 HyperMu transposon, Epicentre) using overlap extension PCR. The overlap extension product was cloned into plasmid pCR8/GW/TOPO (Life Technologies) following manufacturers instructions, and correct sequence was confirmed by Sanger sequencing. These plasmid constructs were then transformed directly into *X. fastidiosa* via electroporation as previously described (38) and transformants were selected by growth on PD3 medium supplemented with chloramphenicol (5 µg/ml) for 10 days at 28°C. Correct transformants were confirmed by PCR amplification and Sanger sequencing of the region of recombination.

## Data availability

The updated Bakersfield-1 assembly has been entered into GenBank under BioProject PRJNA545724. The remaining genome assemblies generated in this manuscript have been entered into GenBank in association with BioProject PRJNA672949. Illumina sequencing reads for Stag’s Leap are available from Jianchi Chen, USDA-ARS San Joaquin Valley Agricultural Sciences Center, on request. All other sequencing reads used for assembly and fast5 files used for methylation analysis have been submitted to Sequence Read Archive (SRA) under BioProject PRJNA672949. *X. fastidiosa* strains sequenced in this study are available from Lindsey Burbank upon request.

## Acknowledgements

Funding for this work was from United States Department of Agriculture (USDA) Agricultural Research Service appropriated project 2034-22000-015-00D. Mention of trade names or commercial products in this publication is solely for the purpose of providing specific information and does not constitute endorsement by USDA. USDA is an equal opportunity provider and employer. We thank Jianchi Chen for providing Illumina sequencing reads for *X. fastidiosa* strain Stag’s Leap.

---

**Figure S1.**
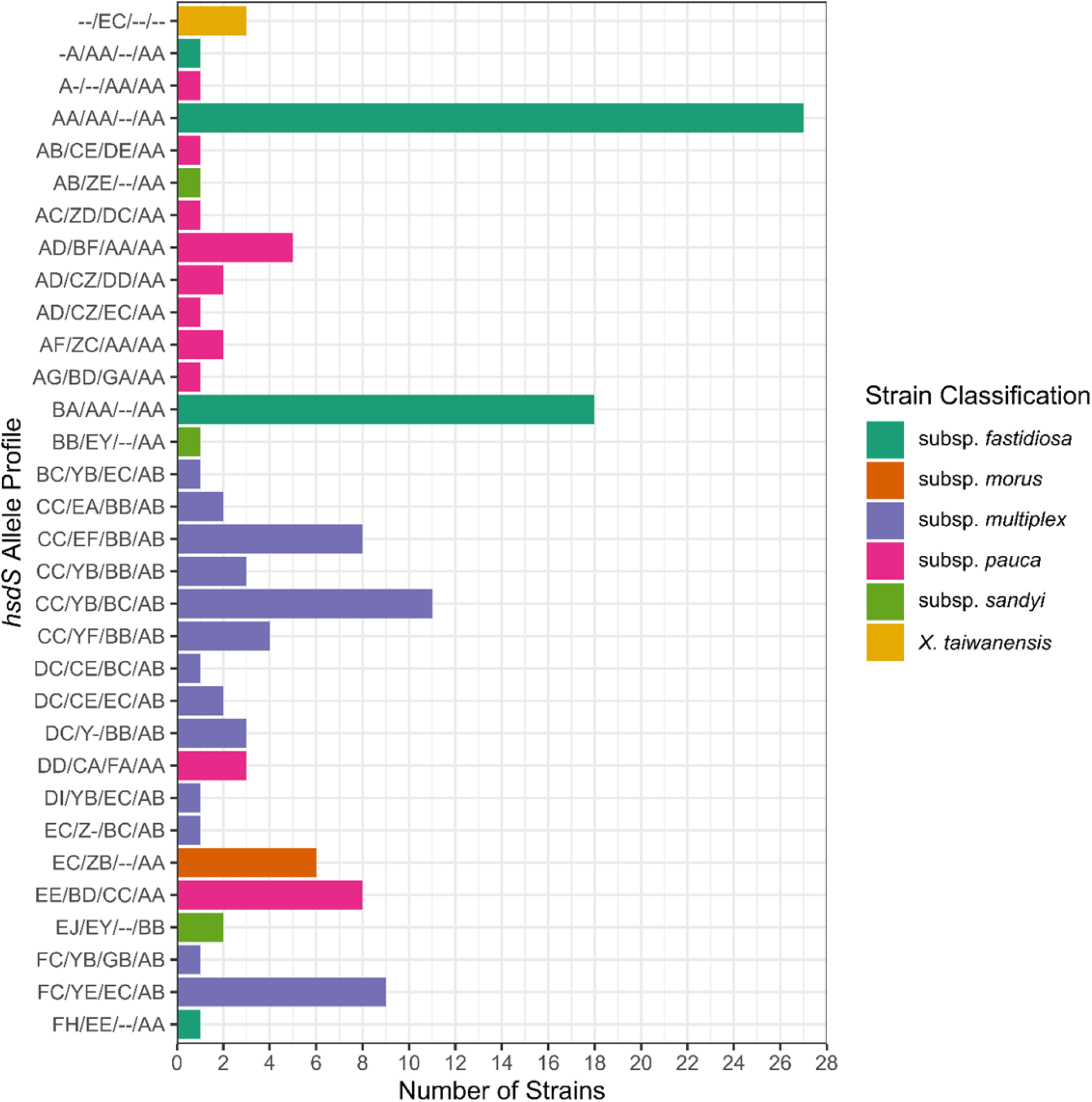
Unique *hsdS* allele profile abundance across *Xylella* strains. Frequency of Type I R-M system *hsdS* allele combinations (e.g., allele profiles) present among *Xylella fastidiosa* subspecies and *X. taiwanensis* strains. The allele profile is presented as a combination of *hsdS* alleles of XfaI/XfaII/XfaIII/XfaIV. “-” indicates no TRD domain was identified for the corresponding TRD position (i.e., TRD1 and/or TRD2). Note that only strains belonging to *X. fastidiosa* subspecies *multiplex* and *pauca* contain XfaIII.

**Figure S2.**
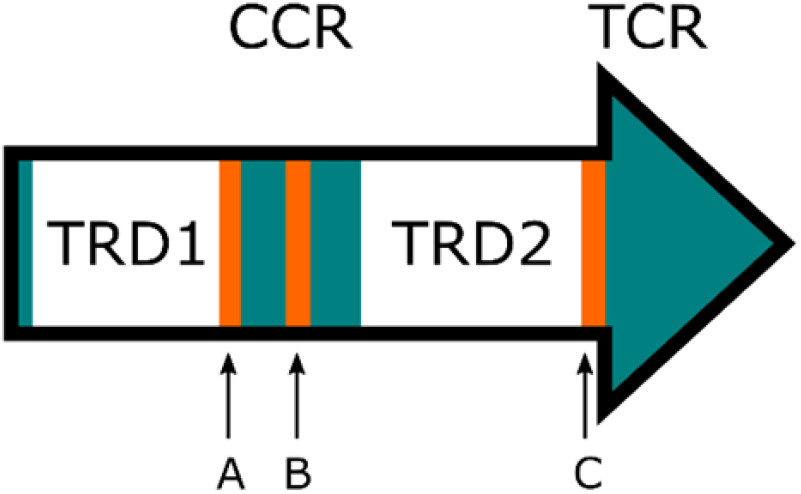
XfaII-*hsdS* contains multiple simple sequence repeat tracts. *HsdS* genes belonging to XfaII contain up to three simple sequence repeat (SSR) tracts, indicated in orange. Conserved regions are indicated in teal. CCR = core conserved region; TCR = terminal conserved region; TRD1 = target recognition domain 1; TRD2 = target recognition domain 2. SSR tracts are labeled as follows: A, tract of 7-9 cytosine bases in TRDs E, Y, and Z; B, the 12 base tract with 0.75 to 11.75 repeats, corresponding to the LEAE tetraamine repeat in *X. fastidiosa* strains; C, a tract of 7-9 cytosine bases in all alleles.

**Table S1**. Type I R-M system components identified in *Xylella* strains in GenBank annotations. “-” indicates that component was not identified in that strain.

**Table S2**. *HsdS* alleles identified in *X. taiwanensis*-specific Type I R-M systems.

**Table S3**. Geographic origin, host, and isolation year of *Xylella* strains used in this study. “*” indicates these strains were isolated from plant material imported from this location. “-” indicates no information was available.

**Table S4**. Additional sequencing and assembly information. “*” indicates this strain was assembled with Unicycler.

